## Supplemental data for "TRPV1 drugs alter core body temperature via central projections of primary afferent sensory neurons"

### Supplementary data

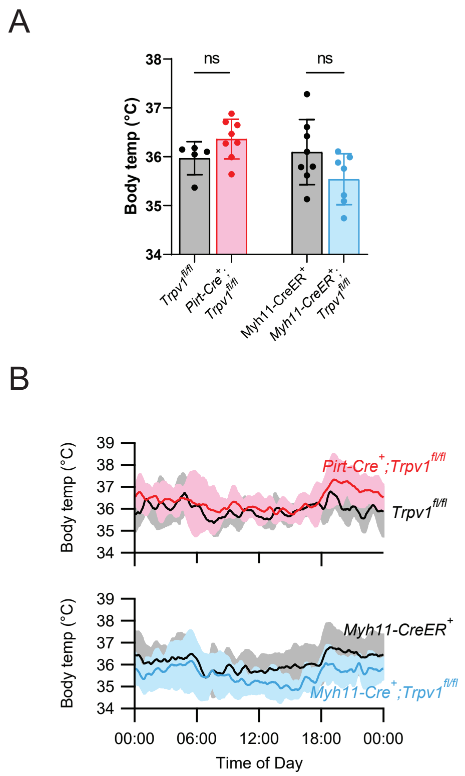


**Figure 1–figure supplement 1** Average core body temperature (**A**) and circadian body temperature fluctuations (**B**) of the indicated genotypes. Unpaired, two-tailed Welch's t-test in (**A**), with each datapoint representing a single mouse. Traces in (**B**) were smoothened by taking running averages (see **Materials and Methods**).

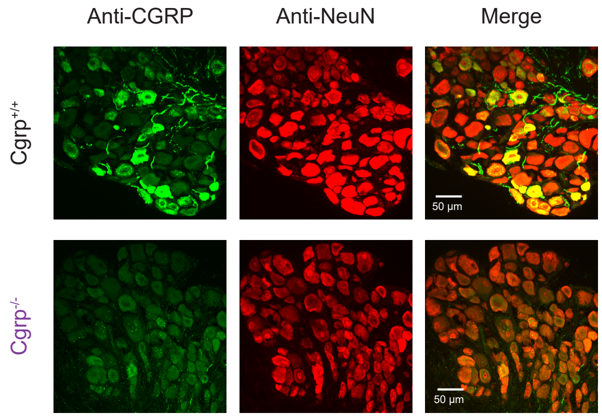


**Figure 3–figure supplement 1** Successful ablation of CGRP expression in *Cgrp*$^{-/-}$ mouse line as shown by the lack of anti-CGRP immunosignal (green) in an example DRG section from a *Cgrp*$^{-/-}$ mouse as compared to a wildtype littermate control. Anti-NeuN (red) is included to reveal all neuronal cell bodies.
